# Cataract-associated deamidations on the surface of γS-crystallin increase protein unfolding and flexibility at distant regions

**DOI:** 10.1101/646083

**Authors:** Heather M. Forsythe, Calvin Vetter, Kayla Ann Jara, Patrick N. Reardon, Larry L. David, Elisar J. Barbar, Kirsten J. Lampi

## Abstract

Deamidation is a major age-related modification in the human lens that is highly prevalent in crystallins isolated from cataractous lenses. However, the mechanism by which deamidation causes proteins to become insoluble is not known, because of only subtle structural changes observed *in vitro*. We have identified Asn14 and Asn76 of γS-crystallin as highly deamidated in insoluble proteins. These sites are on the surface of the N-terminal domain and were mimicked by replacing the Asn with Asp residues. We used heteronuclear NMR spectroscopy to measure their amide hydrogen exchange and ^15^N relaxation dynamics to identify regions with significantly increased dynamics compared to wildtype-γS. Changes in dynamics were localized to the C-terminal domain, particularly to helix and surface loops distant from the mutation sites. Thus, a potential mechanism for γS deamidation-induced insolubilization in cataractous lenses is altered dynamics due to local regions of unfolding and increased flexibility.

## Introduction

Cataracts are the leading cause of low vision over age 40 in the United States and the leading cause of blindness worldwide. The number of individuals affected continues to increase due to our aging population (*1*). Lens transparency relies on the major structural proteins, called crystallins to achieve short range order, compact structures, and close packing at the high concentrations found in the lens (*2*).

The γS-crystallin is in high abundance in human lenses, about 9% of the total crystallins in the young lens (*3*), and shares a high degree of homology within the β/γ-crystallin family. The β/γ-crystallins contain about 35% β-strand content in extremely stable Greek key motifs necessary for their long life spans (*4*), and about 30% disordered content, including the extensions of the N- and C-terminal domains (N-td and C-td) (*5*). The γS crystallin is a ∼21kDa globular protein consisting of four highly stable Greek key motifs joined by disordered loops, with two structurally mirrored halves connected by an unstructured linker (Fig. 1). This monomeric structure solved by NMR (*6, 7*) is consistent with the known C-td (*8*) and dimer (*9*) crystal structures.

**Figure 1.**
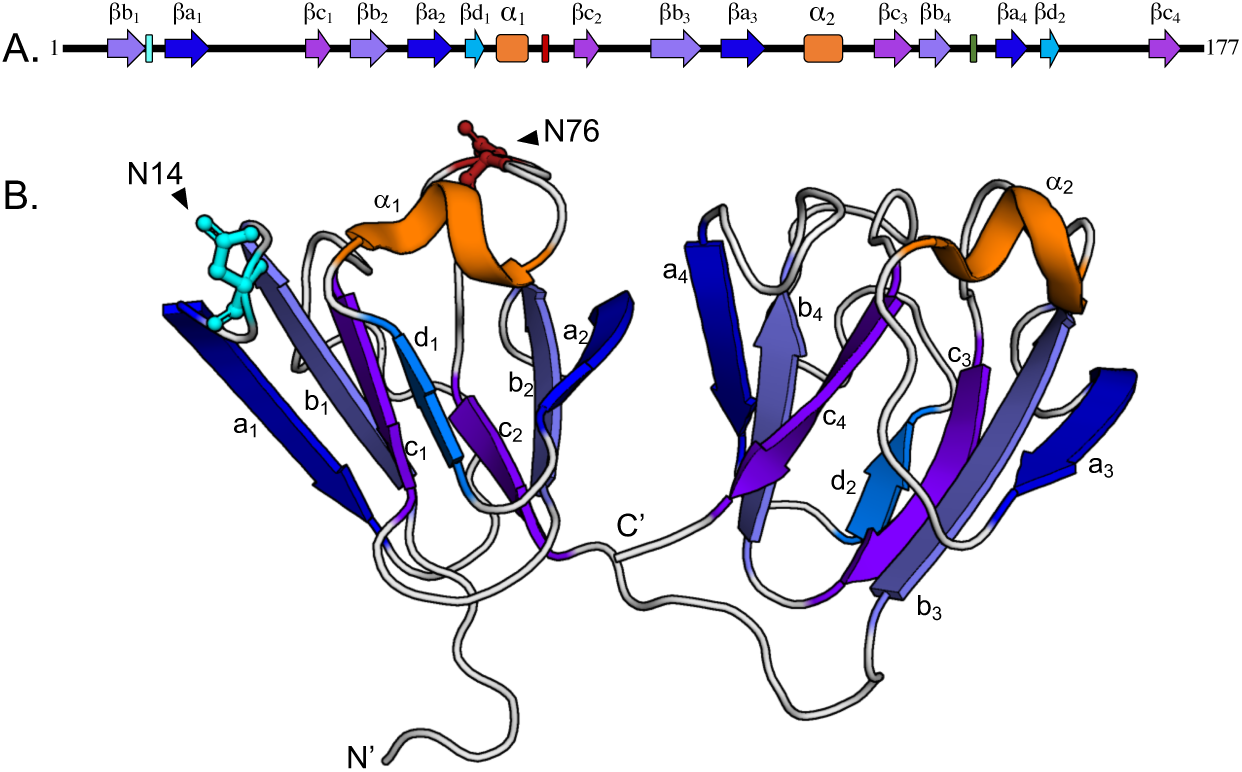
Location of labile Asn residues, N14 and N76, in γS. **(A)** N14 and N76 are in the N-terminal domain between β-strands b_1_ and a_1_ of Greek key motif 1 and between α-helix_1_ and β-strand c_2_ of Greek key motif 2, respectively. (**B)** Ribbon structure showing N14 and N76 located on the surface of the N-terminal domain (PDB: 2M3T).

We have identified deamidation as a major age-related modification in the lens and determined it is associated with insolubilization of crystallins *in vivo* (*10, 11*). Deamidation can occur by nucleophilic attack of the amide nitrogen on the tertiary carbonyl group of an Asn (or Gln) residue resulting in an imide ring and release of the tertiary amino group. When the ring is cleaved in aqueous environments, the resulting Asp (or Glu) carboxylate group introduces a negative charge. Deamidation of Asn *in vivo* leads to isomerization (β-linkage) and epimerization (conversion of l to d stereoisomers) of resulting Asp residues (*12, 13*).

Deamidation occurs in both cataractous and aged clear human lenses (*13-15*) with several sites more deamidated in cataractous lenses compared to age-matched controls (*10, 16, 17*) and at more surface accessible sites (*12, 13*). In normal aged lenses, we identified the highest abundance deamidations in γS in the water-insoluble proteins at Asn 14 (N14) and Asn 76 (N76). The extent of deamidation was roughly twice that found in the water-soluble proteins (*10*). These deamidation sites are well documented in the literature (*12*). In particular, deamidation at N76 in γS has been implicated in cataracts (*13*).

We have mimicked deamidation *in vitro* by replacing Asn/Gln residues with Asp/Glu residues in human β-crystallins, which leads to subtle structural changes and decreased stability (*4, 18-22*). During unfolding, deamidated crystallins rapidly aggregate and are not rescued by α-crystallin chaperone with both proteins precipitating (*18, 19*). Similar effects of deamidation on γD and γS have been observed by others (*13, 23, 24*). In γS, deamidation at N76 leads to decreased stability during chemical unfolding and ineffective chaperoning by α-crystallin (*25*). Deamidation at N76 and Asn143 together led to higher attractive interactions and a tendency towards aggregation with little structural perturbation (*24*). However, none of these studies identified specific residues involved in the altered interactions that led to decreased stability or aggregation.

Here we report that mimicking deamidations at N14 and N76 located on the surface of γS in the N-td disrupt regions in the C-td distant to the deamidation site, exposing a helical region in the C-td and altering surface loop regions that are known protein interactions sites, thus providing a potential mechanism for aberrant interactions of γS and its insolubilization at the high levels in cataractous lenses.

## Results

### Deamidations at Asn14 and Asn76 in γS-crystallin

Human γS-crystallin contains 5 Asn and 9 Gln residues that are potential deamidation sites, of these, N14 and N76 are especially labile and are found on the surface of the N-td (Fig. 1). The N14 is between β-strands b_1_ and a_1_ of Greek key motif 1 and N76 is between α-helix_1_ and β-strand c_2_ of Greek key motif 2 (PDB:2M3T). Due to the presence of the most aged proteins in the nucleus of the lens, water-soluble and water-insoluble proteins were isolated from normal 82-year old and brunescent 85-year old lenses and the level of deamidation at N14 and N76 determined by liquid chromatography/mass spectrometry analysis of tryptic peptides 7-ITFYEDK<u>N</u>FQGR-18 and 72-WMGL<u>N</u>DR-78 (Fig. 2). Deamidation of N14 and N76 was 77 and 72%, respectively, in the water-insoluble fraction of the nucleus of the brunescent lens (Figs. 2a-c and S1). Deamidation at these sites was more abundant in the brunescent lens than the nearly age-matched normal clear lens. Both deamidations were associated with water-insolubility, especially for deamidation at N76, which was 11-fold more abundant in the water-insoluble fraction than the soluble fraction. Deamidation was also detected at Q16, but at only 24% and occurred most often in peptides that also contained deamidation at N14 (Fig. S1). These results suggested that deamidations at N14 and N76 could have contributed to the insolubility of γS-crystallin in the brunescent lens.

**Figure 2.**
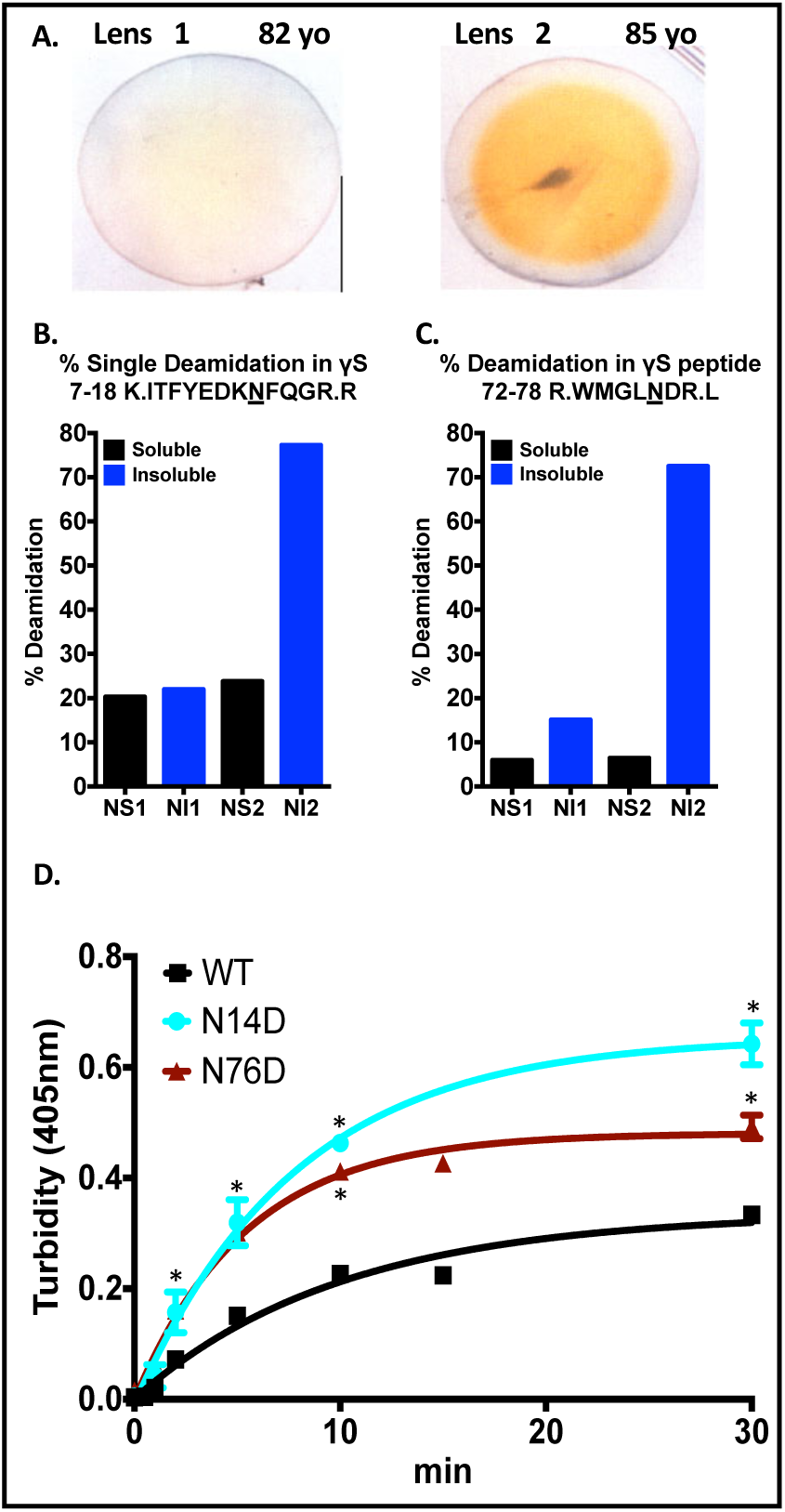
Deamidation at γS N14 and N76 is associated with insolubility in aged lenses and increases susceptibility to heat-induced aggregation. **(A)** Photographs of 82-year old normal (lens 1) and 85-year old brunescent (lens 2) used to measure the extent of deamidation at γS N14 and N76 in the nuclear (center) region of the lens. **(B and C)** The % deamidation at γS N14 and N76, respectively, were measured by mass spectrometry of tryptic peptides from the nuclear soluble (NS) and nuclear insoluble (NI) fractions of lenses 1 and 2 shown in A. **(D)** Relative turbidity of γS-crystallins during thermal-induced denaturation of WT (black), γS N14D (cyan), and γS N76D (red). Proteins at 24 µM were incubated at 70 °C and turbidity measured as change in O.D. at 405 nm, n=3 and SEM calculated via 2-way ANOVA with p < 0.05 (*). Curve fit was generated using nonlinear regression with PRISM 6 (GraphPad Software, CA).

To determine potential changes in γS-crystallin stability and structure due to deamidation at N14 and N76, deamidation was mimicked by site-directed mutagenesis, replacing the Asn with Asp residues at these positions. The protein aggregation of these mutants was determined by measuring their turbidity during thermal-denaturation at 70 °C (Fig. 2d). This temperature was chosen, because it is above the midpoint of the thermal transition for WT Lower temperatures of 60 and 65 °C did not lead to detectable turbidity after 2 h. Aggregation was significantly greater in the N14D and N76D mutants compared to WT after 2 min. The initial rate of increase in turbidity was also significantly greater in both deamidation mimics, 0.06 min^−1^, compared to WT, 0.03 min^−1^ (Student’s t-test, p=0.04) suggesting decreased stability.

### Dynamics of WT γS crystallin determined by NMR

NMR spectra collected on the monomeric γS crystallin showed widely dispersed peaks indicative of a well-ordered protein and consistent with published NMR structures (Fig. S2) (*6, 7*). Resonance assignments of ∼90% of backbone amides aided by previously assigned spectra (*7*) were confirmed by 3D HNCACB spectra collected on ^13^C/^15^N samples. The few missing peaks were for residues in the C-td in βb3, βc3, and βd2, and the flanking loops, similar to what was reported earlier (*8*). Missing peaks could be due to peak broadening arising from internal backbone flexibility or from molecular aggregation. Spectra at 10-fold dilution did not show any additional peaks, suggesting conformational exchange from backbone flexibility at microseconds time scales.

To identify fast exchanging amide protons that are on the protein surface or in flexible loops, we used CLEANEX experiments that measure backbone proton amide (H/H) exchange with water (*27*). Peaks observed in the CLEANEX spectra (Fig. 3a, top row WT and Fig. S2) corresponded to residues in the most exposed regions of the protein. As expected, the most rapidly exchanging residues (the peaks with the highest intensities) were in the N-terminal extension (residues 3-5), loops (residues 27-32, 75, 165-166), and in the linker connecting the N-td and C-td (residues 91-92). These residues also showed multiple conformations in the NMR ensemble structure (PDB: 2M3T) (*6, 7*). Not expected from the structure, was the relatively fast exchange observed for residues 64 and 65 within βd1 and to a lesser extent at the ends of β-strands a1, c1, a2, b2, a3, a4, and c4.

**Figure 3.**
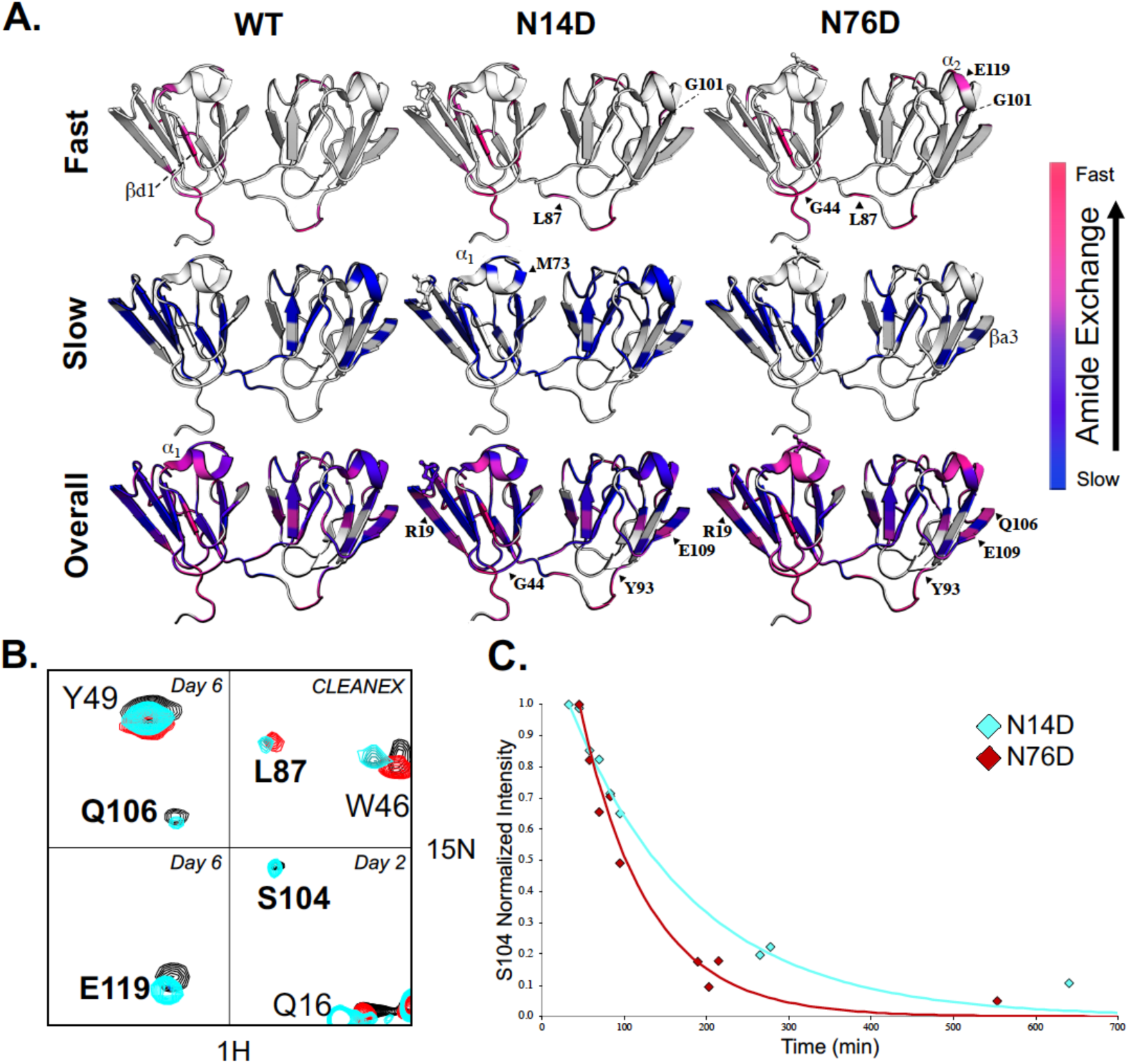
Hydrogen exchange of WT γS-crystallin and N14D and N76D variants measured by NMR. **(A)** (Top) H/H exchange of WT (left), N14D (middle), and N76D (right) showed altered solvent accessibility compared to WT. High intensity peaks that appeared in CLEANEX spectra for amide protons that exchanged the fastest are mapped on the structure in hotpink and lower intensity CLEANEX peaks are mapped in magenta. (Middle) H/D exchange showing the slowest, core residues mapped in blue whose intensity did not significantly change after fourteen days in D_2_ O. (Bottom) Intermediate exchange rates with purple denoting residues that exchanged before the first Best-TROSY spectrum was obtained (in D_2_ O for ∼30min) but weren’t visible in CLEANEX, followed by residues that exchanged after 1 day in D_2_ O (pink), after 2-3 days (magenta-purple), and after 6 days in D_2_ O (dark purple). Those residues that did not have NMR peak assignments available or that were too broad to confidently measure their exchange rates, were mapped on the structure in white. (**B)** NMR spectral overlay of WT (black), N14D (cyan) and N76D (red). Residues which showed significantly altered H/D exchange compared to WT were in bold. Day 6 H/D exchange spectra of Q106 and Y49 (top left), day 6 H/D exchange spectra of E119 (bottom left), CLEANEX spectra of L87 and W46 (top right), and day 2 H/D exchange spectra of S104, Q16, and R145 (bottom right). (**C)** H/D exchange rates of S104 are different for N14D (cyan) and N76D (red) and are reported in Supplementary Table 1.

To determine the slowly exchanging amide protons that are buried and/or involved in H-bonding, we measured hydrogen deuterium (H/D) exchange over a period of two weeks (Fig. 3a, middle row WT). A large number of residues in the β-strands never exchanged significantly (blue) within this time (Figs. 3a, middle row and S2). The interface between the N-td and C-td was not readily solvent accessible (Fig. 3a, magenta-to-purple, bottom row WT). This finding is similar to the slow exchange we have reported for the domain interface in β-crystallins (*5*). Within the flexible connecting peptide, slowly exchanging amide protons were detected for residues 85-87 near the N-td suggesting they were not flexible, but buried within the interface (Fig. 3a, blue, middle row WT). On average, amide protons within the C-td were slower exchanging than those in the N-td. In particular, α_2_ exchanged more slowly than α_1_, as did the connecting loops in the C-td compared to those in the N-td. Exchange rates for amide protons measured in this time window of 1-6 days, and that are shown in Fig. 3a (pink to purple, bottom row WT), are listed in Table S1.

In summary, both H/H and H/D exchange data show that the WT N-td was more dynamic and heterogeneous relative to the C-td, most prominently at N-terminal extension residues and linker residues connecting the domains, except for residues at the N-terminal side that were buried and part of the slowly exchanging protein core. Additionally, H/D exchange data revealed a faster rate of exchange in α1 in the N-td than in α2 in the C-td.

### Effects of mimicking deamidation on γS structure and dynamics

Comparison of WT spectra to those of Asn-to-Asp mutations showed chemical shift differences at the sites of mutation as expected, but also significant differences at unexpected sites distant from the mutation site (Figs. S3 and S4). These sites included residues in the region between βc1 and βb2, α-helix α1, residue L87 in the connecting peptide, C-terminal interface β-strand βa4, α-helix α2, and the most distant β-strand, βa3.

The same or nearby residues observed to have altered chemical shifts in mutant constructs also had significantly altered amide H/H and H/D exchange measurements (Figs. 3a-c and 4a). With N76D, faster exchange rates were observed in general (Figs. 3a and S5). Several new peaks appeared in the CLEANEX spectra, including L87 and G101 in both mutants, as well as G44 and E119 in N76D (Fig. 3a, top row). Peaks for residues newly appearing in the CLEANEX experiment, such as L87, were of greater intensity in N76D than in N14D. Residues with faster exchange rates compared to WT in both mutants included those of βa1 (blue to magenta/pink), and E109 and S104 in βa3 (deep blue to purple) (Figs. 3a, bottom row and 3b, c and Table S1). In N76D, exchange rates were elevated throughout the protein, in particular α_1_ residues (purple to pink) and E119 in helix α_2_ (blue to pink). Furthermore, those residues with increased exchange rates in N14D such as S104 (cyan in overlaid spectra) showed significantly more increase in N76D, disappearing by day 2 (red) (Figs. 3b, c). In both mutants, backbone NH S104 exchanged faster in N76D and N14D than in WT. Conversely, WT NH S104 is part of the slowly exchanging core (Fig. 3a, middle row, near G101 visible in structure).

Compared to WT γS, exchange rates were decreased for some residues in the N14D, but not in the N76D mutant, as shown for the examples M107 in the βa3 (Fig. 3a, magenta to purple, bottom row N14D, near E109), and G44 (Fig. 3a, magenta to purple, bottom row, N14D). Overall, there was a higher number of amide protons that exchanged within 1-6 days measured for N14D and N76D compared to WT (31 for N14D, and 24 for N76D, but only 18 for WT) (Table S1).

To compare dynamics between deamidation mimics and WT γS at the μs to ps time scale, we measured longitudinal and transverse relaxation rates, R_1_and R_2_, and heteronuclear NOE (Figs. 4b and S6). The R_2_/R_1_ values for WT γS, shown in the top row of Fig. 4b, were the most informative in identifying regions of heterogeneous dynamics, because of the clear dynamically heterogeneous values ranging from ∼5-35. Regions with the highest R_2_/R_1_ values included β-strands within the first two Greek key motifs, βa1-4, and βb1-4, and both α-helices, α1 and α2. Those with the lowest R_2_/R_1_ values included the unstructured N-terminal extension and the loop connecting the β-strands βc1, βb2 and βb3. Additionally, linker residues 90-92 had significantly decreased values compared to other residues in the linker.

**Figure 4.**
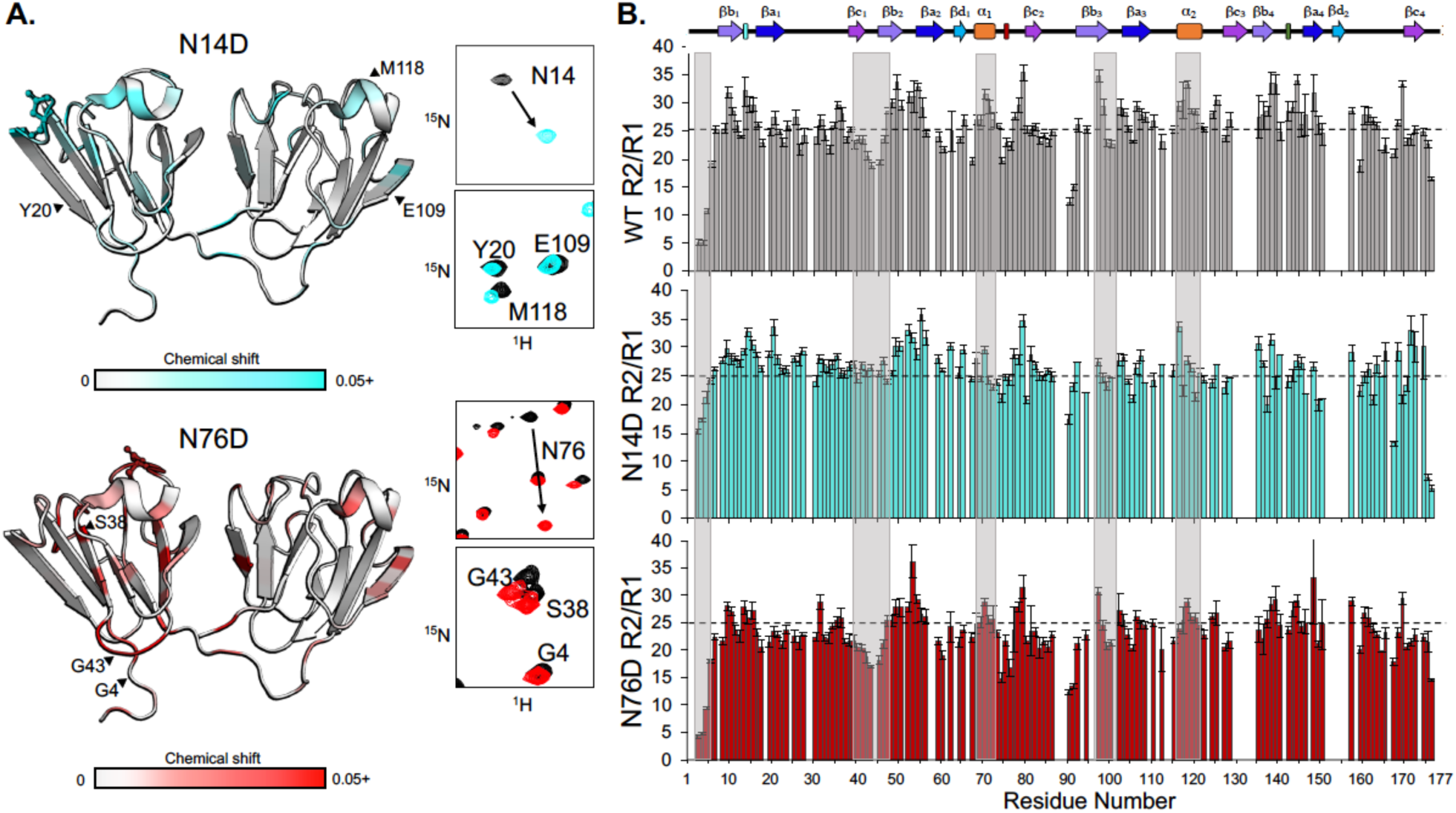
^1^H-^15^N chemical shifts and R_2_/R_1_ values for WT, N14D, and N76D. **(A)** Chemical shift differences mapped on the structure in N14D (top) and N76D (bottom). Shifts shown range from 0.02-0.8 to 0.02-0.15 ppm for N14D and N76D, respectively, and are colored on the structure here from 0-0.5 ppm to distinguish small changes. Full mutant spectra overlaid with WT are shown in Fig. S3, and chemical shifts by residue are presented in Fig. S4. (**B)** R_2_/R_1_ plots of WT (grey), N14D (cyan), and N76D (red). Boxes highlight key differences. The dotted line was placed at the WT average of 25 for comparison.

The R_2_/R_1_ values for N14D showed that there were more homogeneous dynamics in the N-td compared to the C-td, which remained dynamically heterogeneous (Fig. 4b, middle row). For example, residues in the N-terminal extension and in the loop residues between βc1 and βb2 have higher values compared to WT. Conversely, N76D showed similar heterogeneity relative to WT, but with lower values overall (Fig. 4b, bottom row). Heteronuclear NOE measurements for both mutants were on average similar to WT with only modest decrease (Fig. S6). From R_2_/R_1_, we determined rotational correlation times of 11.9, 11.8, and 11.3 ns for WT, N14D, and N76D, respectively. The lower correlation time estimated for N76D suggests that mimicking deamidation at this site causes faster tumbling than expected for a globular protein of ∼21kDa (*7*) suggesting a global increase in flexibility.

Local mobility parameters of backbone amide N-H vectors interpreted using the FAST-ModelFree analysis (*28*) revealed differences in dynamics at both the ms to µs conformational exchange (intermediate exchange, measured as R_ex_) and motions faster than the overall tumbling correlation time on ns to ps timescale (measured as S^2^, where numbers closer to 1 imply restricted motion and those closer to zero imply fully disordered). Residues with a R_ex_ term have heterogenous dynamics specific for these regions. While WT and N76D had 10 and 7 R_ex_ terms respectively, N14D had 37, which was mostly localized to the N-terminus, suggesting deamidation at N14 resulted in multiple conformations specifically at the N-td (Fig. 5a). The C-td also showed regions of exchange but not to the same extent (29 residues with R_ex_ terms in the N-td vs 8 in the C-td).

**Figure 5.**
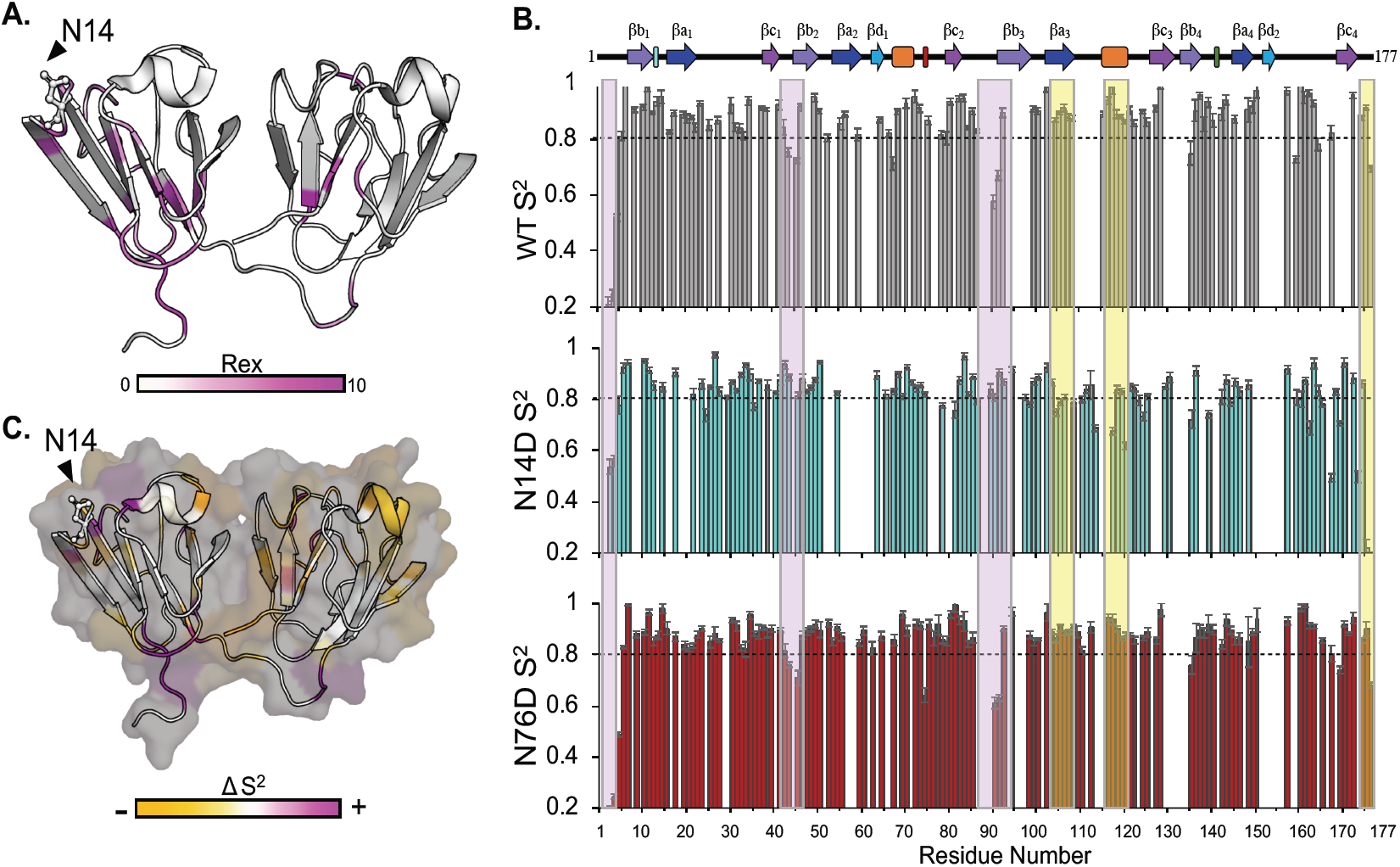
FastModelFree analysis of N14D and N76D. **(A)** N14D R_ex_ values ranging from 0.38-9.01 were mapped on the 3D structure from light to bright magenta. Residues colored white have no R_ex_ valuesdetermined from the model. **(B)** S^2^ plots of WT (grey), N14D (cyan), and N76D (red). Boxes highlight a decrease (yellow) or increase (purple) in S^2^ in N14D. The dotted line at 0.8 was provided for comparison. Few significant differences were observed for N76D at this timescale. **(C)** Changes in S^2^ for N14D were colored by residue onto the structure as a decrease (yellow) or increase in S^2^ (magenta).

Both mutants showed decreased S^2^ values overall, suggesting an increase in fast dynamics. N14D showed significant increases in S^2^ in segments highlighted in purple in the N-td, including in the N-terminal extension, the loop connecting βc1 and βb2, and the end of the linker, leading into the C-td. In contrast, the C-td of N14D showed a decrease in S^2^ specifically in segments βa3, α_2_, and the C-terminal extension, highlighted in yellow (Fig. 5b). Although H/D exchange and chemical shift comparisons show more significant changes in N76D than in N14D, FAST-ModelFree (*28*) analysis revealed more significant changes in N14D.

## Discussion

### Mimicking deamidation causes changes distant from the deamidation site

In both N14D and N76D, the mutation is in the N-td where expected chemical shifts were detected; however, we also detected distant changes in the C-td. Changes in dynamics were observed at the same sites as the chemical shift changes with both N14D and N76D, but at different time scales. With N76D, exchange measurements showed changes in the β-strand, βa3, of the 3^rd^ Greek key motif, and in the helix α_2_ packed against it, suggesting local unfolding specific to this region in the C-td (Fig. 4a). In contrast, changes in this same region in N14D were not detected by hydrogen exchange, but were observed instead as changes in the order parameters obtained from relaxation measurements on the ns to ps time scale. Further changes for N14D R_ex_ were measured with ms to μs fluctuations, primarily in the N-td of N14D reporting multiple conformations.

Changes in dynamics for both N14D and N76D were additionally observed in the connecting linker between domains and at the domain-domain interface itself. The altered dynamics in the N-td likely influence the C-td structure by disruption of these domain-domain interface interactions known to stabilize the overall structure of γS (*29*). We have previously reported similar findings in homologous regions in βB2-crystallin, when the dimer interface was disrupted due to deamidation (*5*). Taken together, these results strongly support that the age-related deamidations on the surface of γS likely contribute to the insolubilization of γS *in vivo* by disrupting multiple regions including the stabilizing interface and those distant from the deamidation site.

A recent study has examined the fast dynamics of WT γS-crystallin and a G57W mutant associated with cataract formation. The G57W mutation perturbed the structure around the mutation site and increased fast dynamics especially in the N-td near the mutation. Substitution of a bulky Trp for the small Gly induced local conformational changes and specifically destabilized the N-td. In contrast, our study examined the impact of deamidation, which does not introduce a large change in side chain size, and measured perturbations at multiple time scales. Importantly, we observed only modest chemical shift perturbations outside of the mutated residues, suggesting that the changes in 3D structure induced by deamidation are small. Yet, dynamic perturbations in the ns to ps and slower motions probed by hydrogen exchange and relaxation measurements detected significant changes occurring in the C-td, despite the mutations being in the N-td.

Our findings of altered protein dynamics at sites distant from the mutation site provide a plausible mechanism for the insolubilization observed *in vivo*. While, preventing age-related deamidation may not be possible, stabilizing the disrupted structural regions might be, such as stabilizing the disrupted interface between domains. A limitation of our findings is that further perturbation of the structure would be expected *in vivo* due to the associated isomerization and epimerization that could not be mimicked here with mutagenesis.

### Deamidation changes highlight potential sites of aggregation

Mimicking deamidation of γS-crystallin at N14 and N76 resulted in altered dynamics at exposed loops both in the N-td and C-td. Several residues that appeared in the CLEANEX experiments conducted on the WT protein are residues previously observed to experience chemical shifts when bound to chaperone αB-crystallin (*6*), particularly G91, S34, E65, and W46, suggesting they are accessible for interactions. In N14D and N76D, residues at the end of the C-td helix, F121 and H122, had increased solvent accessibility. These residues have been reported to interact with αB-crystallin (*6*). Their increased accessibility in the mutants could facilitate interactions with α-crystallin chaperones. And, while initially this might be protective, upon saturation during aging and cataracts, both the γS- and α-crystallins may become insoluble. We have previously reported a similar interaction between α-crystallin and βB2-crystallin (*30*). These surface loops will be explored in subsequent studies for potential sites of interaction.

In summary, residue-specific changes in solvent accessibility and dynamics indicate that surface deamidation sites result in changes both near and distant from the site of deamidation, suggesting that site specific deamidation, has both local and global effects on the protein structure at slow (ms to s) and fast (μs to ps) time scales. The NMR findings combined with the *in vivo* insolubility and *in vitro* aggregation findings support a model that deamidation drives changes that may more readily unfold the protein and expose regions buried in the WT protein. These events in the aged lens could disrupt interactions with other crystallin subunits and increase susceptibility to aggregation, insolubilization, and light scattering (Fig. 6).

**Figure 6.**
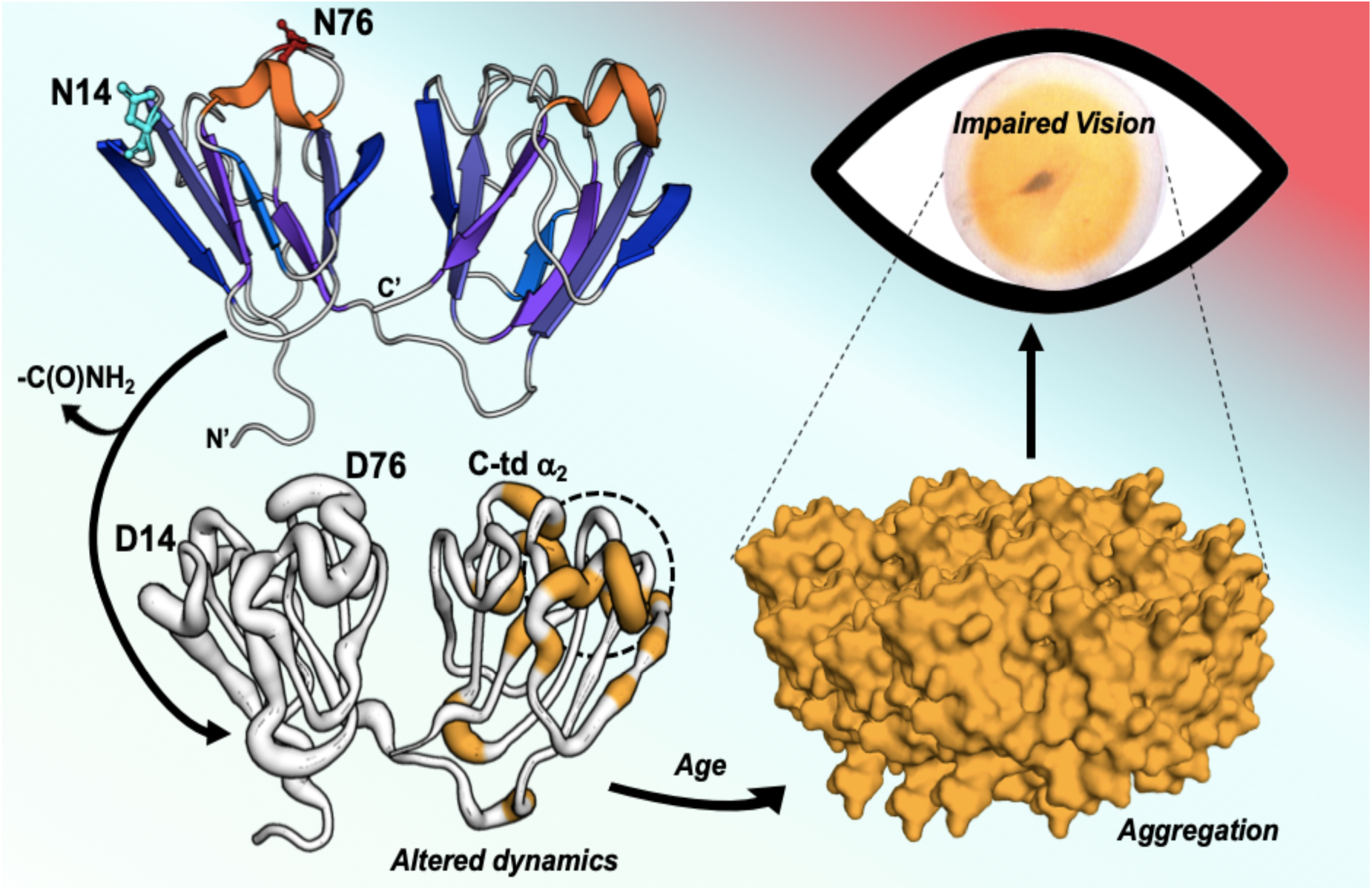
A model showing the effect of deamidation at sites N14 and N76 in γS-crystallin. Increase in dynamics shown as increased thickness in the ribbon structure in the N-td is transferred to localized regions within the C-td. Over time, dynamic changes exposing regions normally buried in the WT protein promoting aberrant interactions with other crystallin subunits and increase susceptibility to aggregation in the aged lens.

## Materials and Methods

### Experimental Design

While, age-related deamidation is thought to be associated with cataracts, the mechanism for deamidation-induced insolubilization of lens crystallins is not known, because *in vitro* studies to date have focused on global changes to protein structure and stability. In this study, we confirmed that extensive deamidation occurs in γS-crystallin at N14 and N76 in the water-insoluble fraction of an aged brunescent lens, mimicked these deamidations using mutagenesis, compared the heat-induced aggregation of these mutants to the WT protein, and measured changes in solvent accessibility and protein dynamics at different time scales using NMR. Use of human donor lenses was approved by Institutional Review Board at Oregon Health & Science University.

### Quantification of modifications at labile Asn in aged lenses

Eyes from donors of 82- and 85-years of age were procured from the Lions Eye Bank of Oregon (approved through the Institutional Review Board at Oregon Health & Science University). Lenses were excised and stored at −70 °C until used. The nuclear region of each frozen lens was isolated using a 4 mm corneal trephine to remove a core from the center of the lens. This core was then homogenized in 20 mM phosphate buffer (pH 7.0) containing 1 mm EDTA, centrifuged at 20,000 x g and the supernatant removed for the water-soluble fraction. The pellet was then washed once and resuspended by sonication for the water-insoluble fraction. The protein content in both fractions was assayed (BCA Assay, Thermo Scientific), lyophilized, and stored frozen at −70°C. Protein samples were resuspended and solubilized in 8M deionized urea, 1.0 M Tris (pH 8.5), 8 mM CaCl_2_, and 0.2 M methylamine. Reduction and alkylation of proteins was then performed using dithiothreitol and iodoacetamide, and samples digested with trypsin (Sequencing grade modified trypsin, Promega, Madison, WI) overnight at 37°C after dilution of the urea to 2M concentration. Digestion was stopped by additional of formic acid, samples centrifuged and the supernatants removed. One 1 µg portions of the digests were injected onto a C18 trap column at 5 μl/min in mobile phase A containing water and 0.1% formic acid, washed for 5 min, then switched on-line to a 2 μm C18, 75 μm x 25 cm EasySpray column (Thermo Scientific) maintained a 40°C with a 300 nl/min flow rate. Peptides were separated by a 7.5-30% mobile phase B (acetonitrile, 0.1% formic acid) gradient over 205 min and analyzed using an EasySpray source and Orbitrap Fusion Tribrid mass spectrometer (Thermo Scientific). Wide isolation single ion monitoring (WiSIM) scans at 500,000 resolution in the Orbitrap to produce high resolution accurate mass data for precursor ions were alternated with data-independent acquisition (DIA) MS/MS scans in the ion trap for sequence confirmation of detected peptides from observed fragment ions. Quantification of the relative amount of deamidation in peptides was determined using Skyline software (*31*), using the 19 mDa mass defect between isotopic peaks of non-deamidated and deamidated peptide ions (*32*). Precursor ion peaks for non-deamidated and deamidated forms of peptides 7-18 (ITFYEDKNFQGR) and 72-78 (WMGLNDR) of γS-crystallin were integrated and % deamidation in each peptide calculated. These calculations included the multiple peptide peaks resulting from isomerization and racemization of N and D residues, as well as the doubly deamidated form of peptide 7-18. The single peak resulting from deamidation at only Q16 was excluded from the calculation of % deamidation at N14, and occurred at a much lower rate than that at N14 as previously reported (*12*). The chromatograms used for these calculations for deamidation at N14 are shown in Fig. S1.

### Recombinant expression of γS-crystallin and deamidated mutants

Deamidation sites were chosen based on the high levels of deamidation and their association with **γ**S-crystallin water-insolubilzation in aged brunescent lenses (Fig. 2). The plasmid containing the gene for human γS WT was engineered into the pE-SUMO (Small Ubiquitin-like Modifier) vector containing a N-terminal 6XHis tagged fusion protein (LifeSensors Inc., Malvern, PA). The γS N14D and γS N76D were generated via site-directed mutagenesis using QuikChangeXL Kit (Agilent Technologies, Santa Clara, CA). *Escherichia coli* cell stocks containing the plasmid of interest were grown at 37 °C, induced for protein expression with IPTG and their growth continued for 4 h. Following cell lysis, proteins were purified using HisPur Cobalt Resin according to the manufacturer’s protocol (Thermo Scientific, Rockford, IL.) and Ulp-1 protease (Ubl-specific protease 1) used to remove the SUMO-tag. Protein was separated from the SUMO-tag by a subsequent pass-through the HisPur Cobalt Resin. Pure protein was aliquoted and stored at −80°C. In all purification steps, 2.5 mM DTT was present to keep the cysteines reduced. Typical yields of protein were several mg per 100 ml of *E. coli* culture. Purity and masses of all expressed proteins were confirmed by mass spectrometry to match predicted masses and be greater than 90% pure. Protein concentration was determined by measuring absorbance of folded protein at 280 nm and calculating concentrations using the protein extinction coefficient calculated using ExPasy ProtParam tool (γS WT 42.860 cm^−1^ M^−1^).

For NMR data collection, γS WT, N14D, and N76D were expressed in MJ9 minimal media at 37°C with ^15^NH_4_Cl as the sole nitrogen source. Additionally, double-labeled WT γS-crystallin was expressed in ^13^C and ^15^NH_4_Cl to confirm assignments. Recombinant protein expression was induced with 0.4 mM IPTG and growth continued at 26 °C for 16 hours. Cells were harvested and purified using TALON His-Tag Purification protocol (Clontech Laboratories, Mountain View, CA.) The SUMO tag was cleaved by Ulp-1 as described above and the protein was further purified using ion-exchange chromatography (Bio-Rad, Hercules, California). In all purification steps, 2.5 mM DTT was present. The purity of the recombinant proteins, as assessed by SDS-polyacrylamide gels, was >95%.

### Heat-induced aggregation of γS-crystallin and deamidated mimics

γS WT was compared to deamidated mimics by measuring changes in their turbidity during heating at 70 °C using previously published methods (*5, 22, 30, 33, 34*). This temperature was chosen based on the midpoint of thermal aggregation at 65°C previously reported (*26*). A 100 μL aliquot of each protein was diluted to 24 μM using 100 mM sodium-phosphate (pH 7.4), 1 mM EDTA, 2.5 mM TCEP, and 100 mM KCl and heated to 70°C in Peltier Thermal Cycler-200 (MJ Research, Hercules, CA). Turbidity of each solution was measured in 96-well plate as the change in optical density at 405 nm using Synergy-HT Microplate Reader (Bio-Tek, Winooski, VT). Data were fit to a nonlinear regression with PRISM 6 (GraphPad Software, CA) and initial rates of aggregate formation determined from the curves. Significance at each time point was determined via 2-way ANOVA with p < 0.05 (*), n=3 with errors bar=SEM.

### Nuclear magnetic resonance of γS-crystallin and deamidated mimics

NMR spectra were collected at 22°C, using 0.5-1 mM ^15^N labeled γS crystallin in a pH 6.9 buffer composed of 10 mM sodium phosphate, 50 mM sodium chloride, 2.5 mM DTT, a protease inhibitor mixture (Roche Applied Science, Madison, WI), and 2-2 dimethylsilapentane-5-sulfonic acid for ^1^H chemical shifts referencing. All NMR experiments were collected on a Bruker 800 MHz Avance III H/D spectrometer equipped with a triple resonance cryoprobe. Assignments previously reported (PDB: 2M3T) were imported from BMRB (accession number 17576). All 2D spectra were processed using Topspin (Bruker Biosciences; RRID:SCR_014227) and analyzed in CcpNmr (*35*). Spectra were referenced to DSS at 0 ppm and chemical shift differences were calculated using the equation δ = [(H shift)^2^ +(N shift/6.51)^2^]^0.5^ (*36*).

### Changes in amide hydrogen solvent accessibility determined by H/H and H/D exchange with NMR

CLEANEX-HSQC experiments for H/H measurements were collected with a mixing time of 100 ms using a recycle delay of 1.5 s. For H/D exchange, NMR samples were dehydrated at room temperature in a Savant SVC 100H Speed Vacuum Concentrator with RH 40-11 rotor and resuspended in an equivalent volume of D2O. We used a Best-TROSY HSQC experiment to assess H/D exchange. The experimental parameters for these experiments were similar to those used in the CLEANEX experiments, except that the recycle delay was reduced to 0.2 s, resulting in a total acquisition time of 13 minutes. The first Best-TROSY spectrum for each sample was taken ∼30min (including the 13min collection time) after resuspension. Six sequential measurements were taken immediately to capture more rapid exchange. Additional measurements were taken over the course of 14 days for a total of 20 time points. Peak intensities were measured as heights determined through CcpNmr Analysis. CLEANEX and core residue peak intensities were normalized to respective TROSY spectra. For non-core resonances in the H/D exchange spectra, the measured intensities were normalized to an average of unaffected residue intensities. H/D exchange rates for non-core resides were calculated by fitting to an exponential decay as described previously (*37*).

### Relaxation measurements and analysis

^1^H-^15^N Heteronuclear NOE experiments were collected with 16 scans, a recycle delay of 8 s, 2048 total points in the direct dimension and 256 total points in the indirect dimension. NOE values were obtained from the ratios of peak intensities in the presence and absence of proton saturation. With I corresponding to peak intensity and δ corresponding to the baseline noise, uncertainty in NOE (σ) was determined using the equation σ/NOE = [(δ_unsat/Iunsat_)^2^ + (δ_sat/Isat_)^2^]^1/2^ (*40*). The delay times for the T_1_ experiment were 0.02, 0.06, 0.1, 0.2, 0.4, 0.6, 0.8, 1.2 s. The delay time for the T_2_ experiment were 16.9, 33.9, 50.9, 67.8, 84.8, 135.7, 169.6, 237.4 ms. All time points were collected in random order and two replicate points were included to assist with error estimation. T_1_ and T_2_ relaxation rates were determined using CcpNmr and error was estimated using the bootstrap method. The T_1_, T_2_ and heteronuclear NOE data were analyzed using five standard models of increasing complexity (*40*). Model selection and model fitting were performed using FastModelFree and ModelFree4 (*28, 38*).

## General

Authors wish to thank Dr. Phillip Wilmarth for careful reading and editing of manuscript.

## Author contributions

Performed all NMR experiments and wrote NMR methods and results (HF-M), Generated recombinant proteins, performed turbidity assays, helped with manuscript preparation (CV), Generated recombinant proteins for NMR (KJ), Operated NMR and helped with manuscript preparation (PR), Performed all mass spectrometry experiments on human donor lenses and wrote methods (LD), Designed all NMR experiments and helped with manuscript preparation (EB), Designed overall experiments and wrote manuscript (KL).

